# Capturing Membrane Trafficking Events During 3D Angiogenic Development in Vitro

**DOI:** 10.1101/2021.04.22.440970

**Authors:** Caitlin R. Francis, Erich J. Kushner

## Abstract

**Objectives:** Mechanisms that govern angiogenesis are paramount to our understanding of how blood vessels are formed embryonically, maintained in adulthood and manifest disease. Akin to transcriptional regulation of endothelial-specific genes, vesicular trafficking events dictate protein localization, functional activity, and half-life, providing a critically important regulatory step. However, there is little information detailing endothelial-specific trafficking signatures. This is due, in part, by limitations in visualizing trafficking events in endothelial tissues. Our aim in this investigation was to explore the use of a 3-dimensional (3D) *in vitro* sprouting model to image and evaluate membrane trafficking events compared to the conventional 2-dimensional (2D)-based culture method.

**Methods:** Primary Human endothelial cells were challenged to make multicellular sprouts using a fibrin-bead assay. An assortment of cell polarity and Rab proteins were quantified via immunocytochemistry and live-imaging to compare their localization between 3D sprouts and 2D culture.

**Results:** Our results show that sprouts generated from the fibrin-bead assay grow close to the imaging plane allowing for an orthogonal view of apical and basal membrane domains. Compared with 2D culture in which the apical and basal domains are in the axial orientation, limiting resolution, 3D sprouts are acquired in the X-Y plane providing high-resolution for viewing trafficking events. Second, we demonstrate that fibrin-bead generated sprouts have a strong apicobasal polarity axis. Third, we directly compare imaging of trafficking mediators podocalyxin and Rab35 between 3D sprouts and 2D culture. Here, we show that 3D sprouting structures are well-suited to capture trafficking events that are not present in 2D culture due to the lack of a defined apical domain. Lastly, we compared exocytic events of von Willebrand Factor between 3D sprouting and 2D culture. Our results demonstrate a distinct imaging advantage for monitoring these trafficking programs in 3D sprouts as compared with conventional 2D culture.

**Conclusions:** In general, our results establish that the fibrin-bead sprouting assay is well-suited for sub-cellular imaging of trafficking events during angiogenic growth. Additionally, the 2D endothelial culture does not enforce the formation of an apicobasal polarity axis.

## INTRODUCTION

The process of early blood vessel formation, or angiogenesis, is critical to establish the requisite vasculature for organismal growth [1]. During angiogenesis, early blood vessels establish the canonical morphologies that define adult vasculature, namely long interconnected conduits providing a portal for blood flow [2]. Endothelial cells are the initial building blocks of blood vessel development, forming small sprouts and capillary-like networks that progressively invade growing embryonic tissue. Our more recent ability to image gross angiogenic processes in both development and disease has unlocked many outstanding questions in the field. This has also been aided by the expanding wealth of transgenic animals as well as a multitude of cellular and molecular biology techniques. However, many processes that are uniquely governed by endothelial tissues and fundamental to their function remain to be explored. In particular, membrane trafficking-based regulation of endothelial function is an underserved area of exploration in the field of angiogenesis that is largely dependent on sub-cellular imaging techniques to interrogate function.

Trafficking broadly refers to the vesicle-based transport and movement of proteins through the cell. This process is parsed into endocytic (outside material in), exocytic (inside material out), and recycling (moving between both endocytic and exocytic) pathways [3-5]. Rab GTPase proteins are the most recognized multifunctional mediators of all trafficking events acting as an intracellular barcoding system with over 70 known family members [6, 7]. Rab proteins attach to the outside of vesicles where they define intracellular trafficking routes [8, 9]. Activated Rabs then recruit or bind to tissue-specific effectors which can then promote localization, signaling, and degradation of their cargo. For example, in ECs Rab27a is required for Weibel-Palade body (WPB) apical membrane fusion and von Willebrand Factor (vWF) exocytosis [10]. Indeed, the breadth of function trafficking encompasses is far and wide, and may collectively represent the most dominant, yet esoteric, regulatory program in a protein’s life cycle [7, 11, 12]. Taken as a whole, how endothelial tissues harness trafficking-based regulation is a major outstanding question in the field of blood vessel development and homeostasis.

Much of the seminal work in the field of membrane trafficking has been carried out in epithelial tissue [13]. This is in large part due to their large rectangular shape and spatially segregated apical and basal domains allowing for relatively easy imaging of processes at either membrane. Additionally, epithelial cells readily set up apicobasal polarity in 2D culture, thus do not require much in the way of physical or chemical cues to elicit a defined polarity axis [14]. By contrast endothelial cells are exceedingly flat exhibiting a mesenchymal morphology [15]. In some instances, the distance between the apical and basal domains in endothelial cells are diffraction limited (≤500nm) hindering imaging of either membrane surface. In 2D culture endothelial cells are highly migratory, setting up a defined planar cell polarity axis; however, removed from a sprouting structure endothelial cells on a dish do not show a commitment to an apical and basal membrane identity. Because trafficking biology primarily entails movement of vesicular cargo, transcript levels of trafficking-mediators are typically held at a constant, excluding the use of simple expression analysis to interrogate function. In this realm, the ability to visualize differential trafficking events to various membranes is paramount to understanding function. Unfortunately, *in vivo* imaging of endothelial trafficking events present a significant challenge, due to the relative thinness of endothelial cells coupled with the requirement for a less magnified microscope objective (e.g. 20x vs 60x lens) to span the tissue depth needed to capture blood vessels. Therefore, we believe the use of a 3D sprouting model is an excellent compromise between providing the necessary cellular cues to reproduce angiogenic morphodynamics with ample sub-cellular imaging accessibility to image trafficking events.

In this investigation our aim was to explore the use of a 3-dimensional (3D) *in vitro* sprouting model to image and evaluate membrane trafficking events in two scenarios. First, our goal was to capture sub-cellular dynamics of apical proteins, in particular Rab GTPases and their cargo, related to lumen biogenesis. Second, was to evaluate luminal exocytic events at the apical membrane. Our reasoning for choosing these trafficking events was two-fold: 1) these events are hard-to-impossible to distinguish in most *in vivo* models; and 2) are typically evaluated using 2-dimensional (2D) culture. Using a fibrin-bead sprouting assay we demonstrate the utility of this 3D *in vitro* system for capturing endothelial-specific trafficking events. Methodologically, we demonstrate that endothelial sprouts develop parallel and close to the imaging window allowing for use of commonly equipped high-resolution objectives. Additionally, we show that lumen biogenesis and exocytic trafficking events are easily imaged in multicellular sprouts to a much greater extent than 2D culture. Overall, this work highlights a highly reproducible *in vitro* assay that provides a tailored imaging platform for exploring blood vessel-specific trafficking networks.

## MATERIALS AND METHODS

A list of used materials is included in the Supplemental Data. The authors will make their raw data, analytic methods, and study materials available to other researchers upon written request.

### Cell culture

Pooled Human umbilical vein endothelial cells (HUVECs) were purchased from PromoCell and cultured in EGM-2 media (PromoCell Growth Medium, ready-to-use) for 2-5 passages. For experiments glass-bottomed imaging dishes were exposed to deep UV light for 6 minutes and coated with Poly-D-Lysine (ThermoFisher) for a minimum of 20 minutes. Small interfering RNA (ThermoFisher) was introduced into primary HUVEC using the Neon® transfection system (ThermoFisher). Scramble and Rab27a siRNAs were purchased from (ThermoFisher) and resuspended to a 10µM stock concentration and used at 0.5 µM (*see data supplement*). Normal human lung fibroblasts (NHLFs, Lonza) and HEK-A (ThermoFisher) were maintained in Dulbeccos Modified Medium (DMEM) supplemented with 10% fetal bovine serum and pen/strep antibiotics. Both NHLFs and HEKs were used up to 15 passages. All cells were maintained in a humidified incubator at 37°C and 5% CO_2_.

### Sprouting Angiogenesis Assay

Fibrin-bead assay was performed as originally reported by Nakatsu et al. 2007 [16]. Briefly, HUVECs were coated onto microcarrier beads (Amersham) and plated overnight. SiRNA-treatment or viral transduction was performed the same day the beads were coated. The following day, the EC-covered microbeads were embedded in a fibrin matrix. Once the clot was formed, media was overlaid along with 100,000 NHLFs. Media was changed daily along with monitoring of sprout development.

### Plasmid Constructs

The following constructs were procured for the study: GFP-Rab27A (gift from William Gahl; Addgene plasmid #89237); Rab35 (gift from Peter McPherson; Addgene plasmid #47424); Neo DEST (705-1) (gift from Eric Campeau & Paul Kaufman; Addgene plasmid #17392); pMDLg/pRRE (gift from Didier Trono; Addgene plasmid # 12251); pVSVG (gift from Bob Weinberg; Addgene plasmid #8454); psPAX2 (gift from Didier Trono; Addgene plasmid #12260); pShuttle-CMV (gift from Bert Vogelstein; Addgene plasmid #16403); and AdEasier-1 cells (gift from Bert Vogelstein; Addgene, #16399)

### Lentivirus and Adenovirus Generation

Lentivirus was generated by using the LR Gateway Cloning method [17]. Genes of interest and fluorescent proteins were isolated and incorporated into a pME backbone via Gibson reaction [18]. Following confirmation of the plasmid by sequencing the pME entry plasmid was mixed with the destination vector and LR Clonase. The destination vector used in this study was pLenti CMV Neo DEST (705-1). Once validated, the destination plasmids were transfected with the three required viral protein plasmids: pMDLg/pRRE, pVSVG and psPAX2 into HEK 293 cells. The transfected HEKs had media changed 4 hours post transfection and the viral media was harvested at day 3.

Adenoviral constructs and viral particles were created using the Adeasy viral cloning protocol [19]. Briefly, transgenes were cloned into a pShuttle-CMV plasmid via Gibson Assembly. PShuttle-CMV plasmids were then digested overnight with MssI (ThermoFisher) and linearized pShuttle-CMV plasmids were transformed into the final viral backbone using electrocompetent AdEasier-1 cells. Successful incorporation of pShuttle-CMV construct into AdEasier-1 cells were confirmed via digestion with PacI (ThermoFisher). 5000 ng plasmid was then digested at 37°C overnight, then 85°C for 10 minutes and transfected in a 3:1 polyethylenimine (PEI, Sigma):DNA ratio into 70% confluent HEK 293A cells (ThermoFisher) in a T-25 flask.

Over the course of 2-4 weeks, fluorescent cells became swollen and budded off the plate. Once approximately 70% of the cells had lifted off the plate, cells were scraped off and centrifuged at 2000 rpm for 5 minutes in a 15 mL conical tube. The supernatant was aspirated, and cells were resuspended in 1 mL PBS. Cells were then lysed by 3 consecutive quick freeze-thaw cycles in liquid nitrogen, spun down for 5 minutes at 2000 rpm, and supernatant was added to 70% confluent T-75 flasks. Propagation continued and collection repeated for infection of 10-15cm dishes. After collection and 4 freeze thaw cycles of virus collected from 10-15cm dishes, 8 mL viral supernatant was collected and combined with 4.4 g CsCl (Sigma) in 10 mL PBS. Solution was overlaid with mineral oil and spun at 32,000 rpm at 10°C for 18 hours. Viral fraction was collected with a syringe and stored in a 1:1 ratio with a storage buffer containing 10 mM Tris, pH 8.0, 100 mM NaCl, 0.1 percent BSA, and 50% glycerol. HUVEC were treated with virus for 16 hours at a 1/10,000 final dilution in all cell culture experiments.

### Quantification of trafficking proteins

Quantification of pHlourin-podocalyxin was determined by projecting images (both 2D and 3D cells/sprouts) in 3D using the FIJI XXX function and visually identifying accumulation(s) of pHlourin-podocalyxin relative to various plasma membranes. Quantification of WPB number was performed by counting vWF puncta on a per cell basis in both 2D culture and 3D sprouts. In both culture conditions, WPB localization was determined by first identifying WPB accumulations (≥4 WPBs) or where the highest population resided in each individual cell. Second, images were rendered as a 3D projection as mentioned above and the WPB puncta was scored based on its most proximal membrane (e.g. basal or apical). Secretion of vWF was scored visually by identifying whether vWF was in a lumen or contained within a cell. Actin stain determined cell and lumen boundaries.

### Immunofluorescence and Microscopy

Prior to seeding cells, coverslips were treated with poly-D Lysine for approximately 20 minutes and washed 2 times with PBS. HUVECs were fixed with 4% PFA for 7 minutes. ECs were then washed three times with PBS and permeabilized with 0.5% Triton-X (Sigma) for 10 minutes. After permeabilization, cells were washed three times with PBS. ECs were then blocked with 2% bovine serum albumin (BSA) for 30 minutes. Once blocked, primary antibodies were incubated for approximately 4-24 hours. Thereafter, primary antibodies were removed, and the cells were washed 3 times with PBS. Secondary antibodies with 2% BSA were added and incubated for approximately 1-2 hours, washed 3 times with PBS and mounted on a slide for imaging.

For imaging the fibrin-bead assay, fibroblasts were removed from the clot with a 1-minute trypsin incubation. Following incubation, the trypsin was neutralized with DMEM containing 10% BSA, washed 3 times with PBS, and fixed using 4% paraformaldehyde for 40 minutes. After fixation, the clot was washed 3 times with PBS, permeabilized with 0.5% Triton-X for 2 hours and then blocked with 2% BSA for 1 hour prior to overnight incubation with primary antibodies. The following day, primary antibodies were removed, and the clot was washed 5 times with PBS and secondary antibody was added with 2% BSA and incubated overnight. Before imaging, the clot was washed 5 times with PBS. All primary and secondary antibodies are listed in the *supplemental data*. Images were taken on a Nikon Eclipse Ti inverted microscope equipped with a CSU-X1 Yokogawa spinning disk field scanning confocal system and a Hamamatusu EM-CCD digital camera. Images were captured using a Nikon Plan Apo 60x NA 1.40 oil objective using Olympus type F immersion oil NA 1.518. All images were processed using ImageJ (FIJI).

### Statistical Analysis

Experiments were repeated a minimum of three times. Statistical analysis and graphing was performed using GraphPad Prism. Statistical significance was assessed with a Student’s unpaired t-test for a two-group comparison. Statistical significance set a priori at p<0.05.

## RESULTS

### Generating 3-dimensional sprouts using the fibrin-bead assay

To image endothelial-specific trafficking signatures we employed a fibrin-bead sprouting assay first described by Nakatsu et al. [20]. In this assay endothelial cells are coated onto a micro-carrier bead and then embedded into a fibrin matrix. Following the addition of a fibroblast feeder layer, the endothelial cells sprout into the surrounding matrix (**Fig. 1A**). Importantly, these sprouts produce multicellular proto-vessel structures, as opposed to solely filopodia invasion such as those observed in the Matrigel assay [21]. Here, sprouts reproduce characteristic *in vivo* sprouting features such as branching, dynamic cell shuffling [22, 23], anastomosis [24], and lumen formation [20] (**Fig. 1B**). There are other 3D sprouting assays that may demonstrate comparable sprouting characteristics; however, in our hands, the fibrin-bead assay produced very distinct multicellular sprouts with a clearly defined tip and stalk cell morphology. This is important, as other assays can invade the surrounding matrix fashioning a cavernous lumen-like cavity, but this type of morphology can be primarily attributed to cyst formation, where the cells breakdown the matrix, but lack canonical sprouting characteristics [25]. A possible drawback to this *in vitro* system is the lack of blood flow-based morphodynamic cell rearrangements. Several groups have published microfluidic devices that incorporate fluid-flow that could be potentially substituted here [26, 27]. Although, given the relative ease and low-cost of the fibrin-bead assay, it is likely more accessible to the average laboratory. Additionally, many developmental trafficking events, such as those in lumen biogenesis, precede blood flow, thus a ‘flow-less’ model would be appropriate in these circumstances.

**Figure 1.**
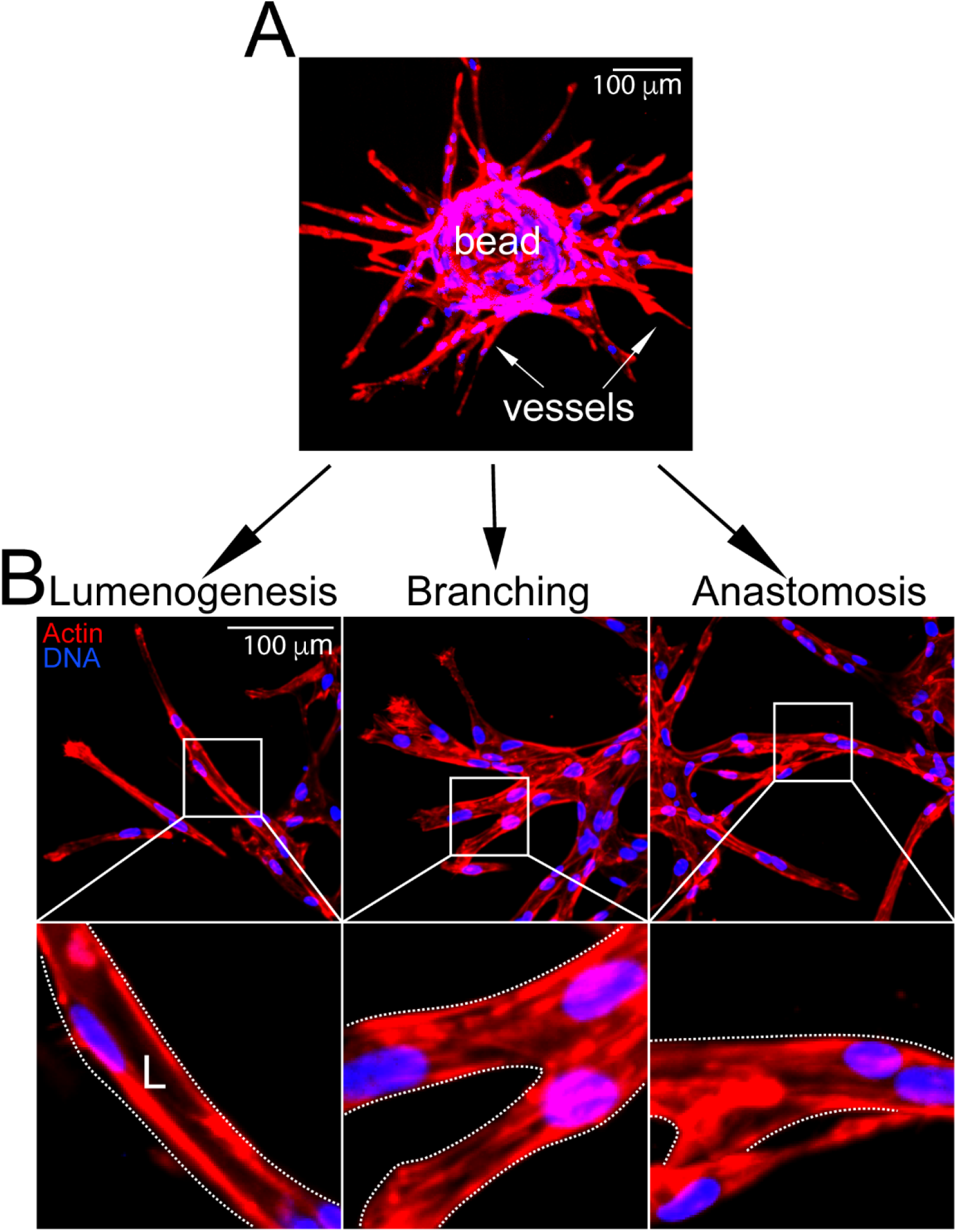
Fibrin-bead assay recapitulates angiogenic traits in vitro. *Top*-representative image of embedded fibrin-bead after 4 days of growth. *Bottom*-representative images of lumenogenesis, branching and anastomosis with magnifications (boxes). L denotes lumen.

### 3D sprouts demonstrate a defined apical and basal polarity

To determine the suitability of the fibrin-bead assay for trafficking studies we first examined if sprout structures demonstrated apicobasal polarity as compared with 2D culture using standard confocal imaging techniques. To test this, 3D sprouts generated in the fibrin-bead assay and endothelial cells plated on coverslips (2D culture) were stained for actin (cytoskeleton), moesin (apical), beta-1 integrin (basal), VE-cadherin (junctional) and podocalyxin (apical). 2D cultured cells demonstrated a diffuse distribution of moesin and podocalyxin with no clear plasma membrane enrichment, indicative of lack of apicobasal polarity (**Fig. 2A**). Orthogonal projections could not provide useful information as the axial resolution was far too low to make out individual puncta; therefore, we did not rely on this method going forward. In 2D, Beta-1 integrin was localized to focal adhesions on the basal surface of the cell (**Fig. 2A**). One interesting note is that endothelial cells cultured on non-compliant surfaces such as hard plastic or glass spread out to a greater extent than cells cultured on soft matrices [28]. This elevated cell spreading in 2D could also contribute to the decrease separation between apical and basal surfaces, complicating discrete imaging of these membrane domains.

**Figure 2.**
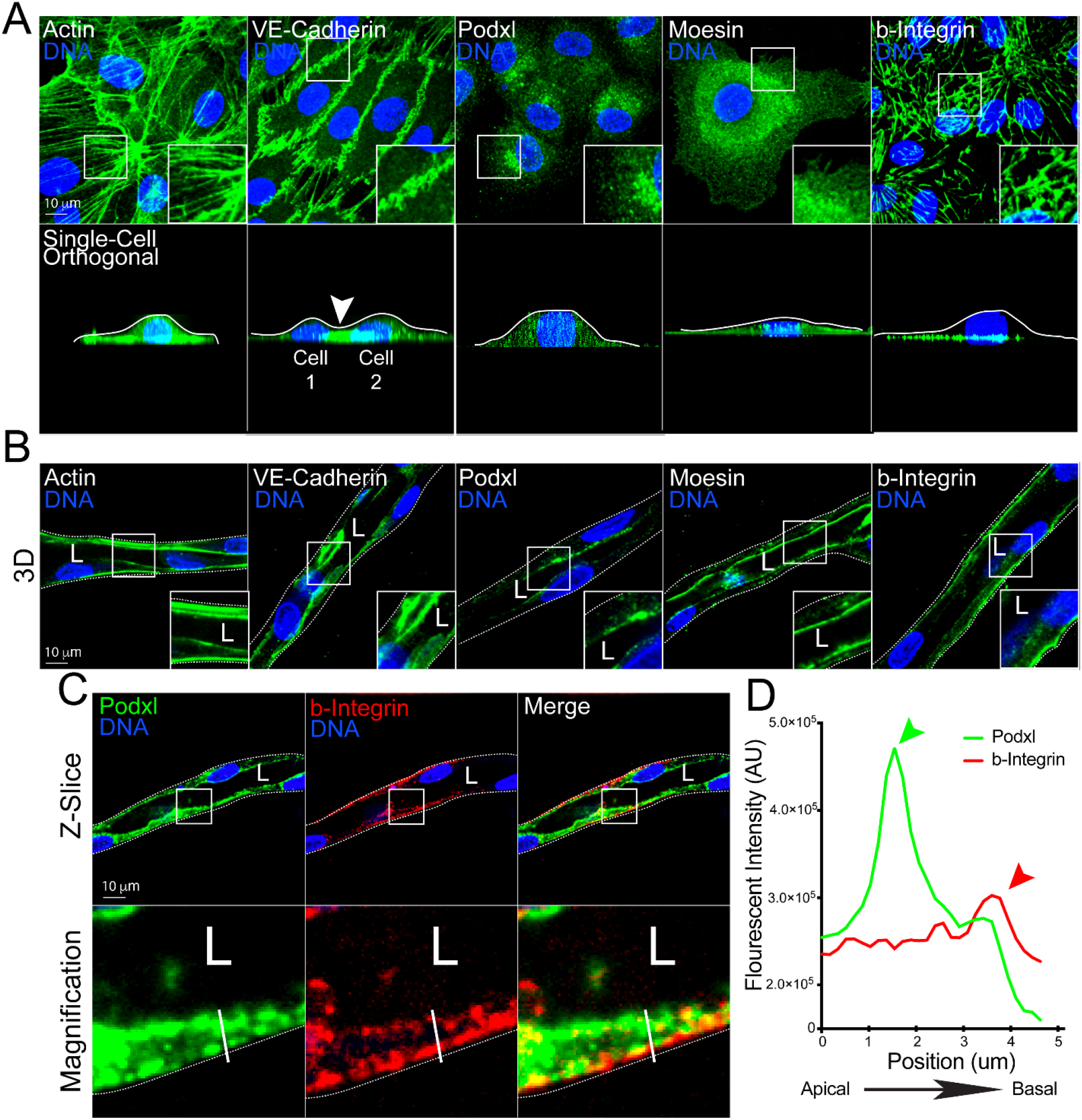
Comparison of cell polarity markers in 2D culture and 3D sprouts. (**A**) Representative images of endothelial cell cultured on 2D surface (top panels) and axial view (x-z plane, bottom panels). Cells were stained with actin (cytoskeleton), VE-cadherin (cell junctions), podocalyxin (Podxl, apical membrane), moesin (apical membrane), and beta-1 integrin (b-integrin, basal membrane). White lines mark apical surface and arrowhead denotes junction between two cells. (**B**) Representative images of fibrin-bead generated sprouts stained for indicated proteins. (**C**) Representative imaging showing co-staining of Podxl and b-integrin within the same sprout cross-section to highlight differences in apical and basal domains. (**D**) Line scan illustrating peaks in fluorescent intensity of podxl and b-Integrin relative to apical and basal domains. The green arrowhead denotes peak of podxl and red arrowhead denotes peak of b-Integrin. The white line in panel C denotes the line scan area. White boxes are areas of magnification and while dotted lines indicated sprout boundaries. L denotes lumen.

An immediately apparent advantage of the fibrin-bead system was that sprouts are oriented in such a manner to the imaging plane that the apical and basal membranes are captured in the X-Y plane as opposed to 2D cells, where the apical and basal domains are in the X-Z orientation (**Fig. 2B**). The X-Z axial plane has significantly lower resolution as compared to images acquired in the X-Y plane. Using a conventional oil 1.4 NA 60x objective with a working distance of 0.21 mm, we encountered no issues imaging sprouts near the coverslip.

In contrast to 2D culture, fibrin-bead generated sprouts exhibited clearly polarized apical and basal membrane surfaces. Sprouts stained for moesin showed a defined accumulation at the apical membrane, distinct from VE-cadherin at cell-cell junctions and beta-1 integrin on the basal membrane (**Fig. 2B**). Similarly, podocaylxin was highly enriched at the apical membrane (**Fig. 2B-D**). We also observed that endothelial cells in sprouting structures displayed increased apical-basal membrane separation as compared with cells cultured in 2D, this also enhanced our ability to distinguish these domains (**Fig. 2C,D**). Overall, these results show fibrin-bead generated sprouts demonstrate apical and basal signaling that can be imaged at high-resolution.

### Dynamic Imaging of podocaylxin trafficking between 2D culture and 3D sprouting

Podocalyxin is a glycoprotein that has been shown to be involved in lumen formation across many developmental models [14]. During lumen formation, podocalyxin is trafficked from the basal membrane to the apical membrane where it orchestrates delamination of the opposing cell membranes, creating the early lumen cavity [29-32]. Although many epithelial studies have documented podocalyxin’s transcytosis, no endothelial studies have live-imaged this trafficking event to our knowledge. To image podocalyxin’s insertion into the apical membrane we constructed a pHluorin-tagged podocalyxin adenovirus (**Fig. 3A**). PHluorin is a green fluorescent protein (GFP) variant that is non-fluorescent in acidified vesicles, but fluorescence is rescued at neutral pH following plasma membrane fusion (**Fig. 3A**) [33]. This approach allows us to differentiate between Podxl that is inserted into the plasma membrane from populations that reside in sub-apical vesicles. We first live-imaged the pHluroin-podocalyxin construct in 2D cultured endothelial cells. Here, podocalyxin was localized to discrete puncta on both the apical and basal domains of the cell (**Fig. 3B,C; Movie 1**); again, suggesting a lack of polarity. By contrast, live-imaging of pHluroin-podocalyxin in fibrin-bead assay sprouts demonstrated a robust and continuous accumulation of podocalyxin at the apical membrane with an elevated localization to sites of lumen deadhesion (**Fig. 3B,D; Movie 2**). In line with previous reports, our results demonstrate that podocalyxin is actively trafficked to the apical membrane; however, it may be trafficked more robustly to cell-cell interfaces actively undergoing membrane deadhesion via undescribed mechanisms. These data also indicate that sub-cellular podocalyxin trafficking events can be captured at high-resolution in the fibrin-bead sprouting assay.

**Figure 3.**
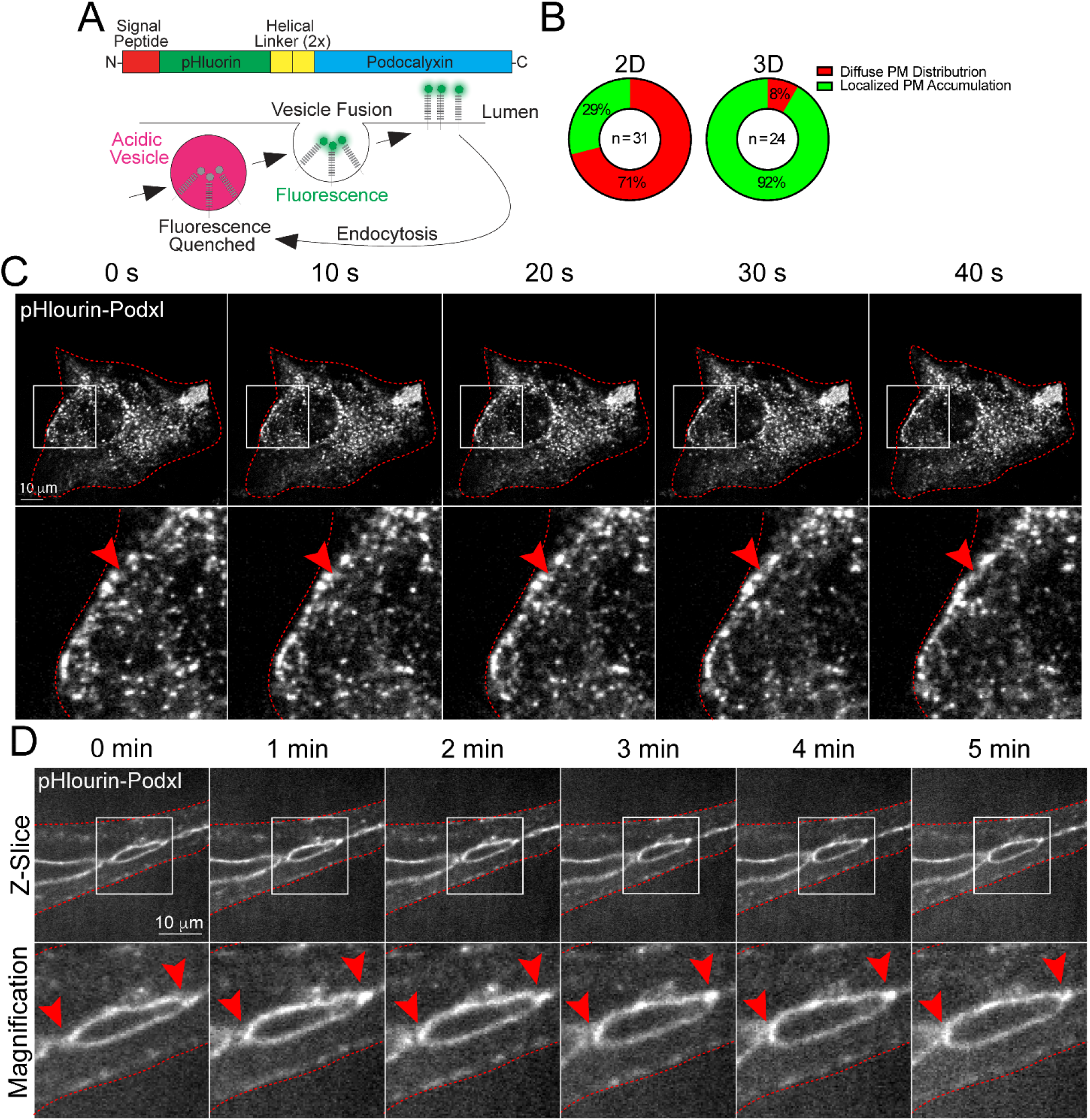
Live-imaging of podocalyxin trafficking in 2D culture and 3D sprouts. (**A**) Structure of engineered pHluorin-Podxl fusion protein. In acidified vesicles, pHluorin fluorescence is significantly quenched. However, once inserted on the plasma membrane at neutral pH, fluorescence is rescued allowing for visualization of plasma membrane insertion. (**B**) Graph showing percentages of pHluorin-Podxl with either even or punctate plasma membrane (PM) distribution. (**C**) Live-imaging of 2D cell expressing pHluorin-Podxl over time. Red arrowheads denote puncta accumulated at the leading edge of the cell. (**D**) Live-imaging of fibrin-bead generated sprout expressing pHluorin-Podxl over time. Red arrowheads indicate active areas of lumen expansion where Podxl is accumulating. White boxes are areas of magnification and red lines indicated sprout boundaries. L denotes lumen.

### Characterizing Rab35 trafficking between 2D culture and 3D sprouting

Rab GTPases represent a class of well-studied proteins that orchestrate vesicle trafficking [6, 34, 35]. In particular, Rab35 has been shown to play a multitude of roles depending on the organism, tissue-type and cellular conditions [36-39]. Also, Rab35’s localization has not been characterized in vascular tissue. Thus, to explore the use of the fibrin-bead assay for imaging endothelial trafficking events, we over-expressed Rab35 in both 2D and 3D culture conditions. In 2D culture, GFP-Rab35 displayed a membranous localization but was broadly distributed, not co-localizing with basal marker Beta-1 integrin, nor cytoskeletal protein actin (**Fig. 4A,B**). By contrast, expression of GFP-Rab35 in 3D fibrin-bead sprouts demonstrated a preference for the apical membrane co-localizing with actin and distinct from beta1 integrin (**Fig. 4C,D**).

**Figure 4.**
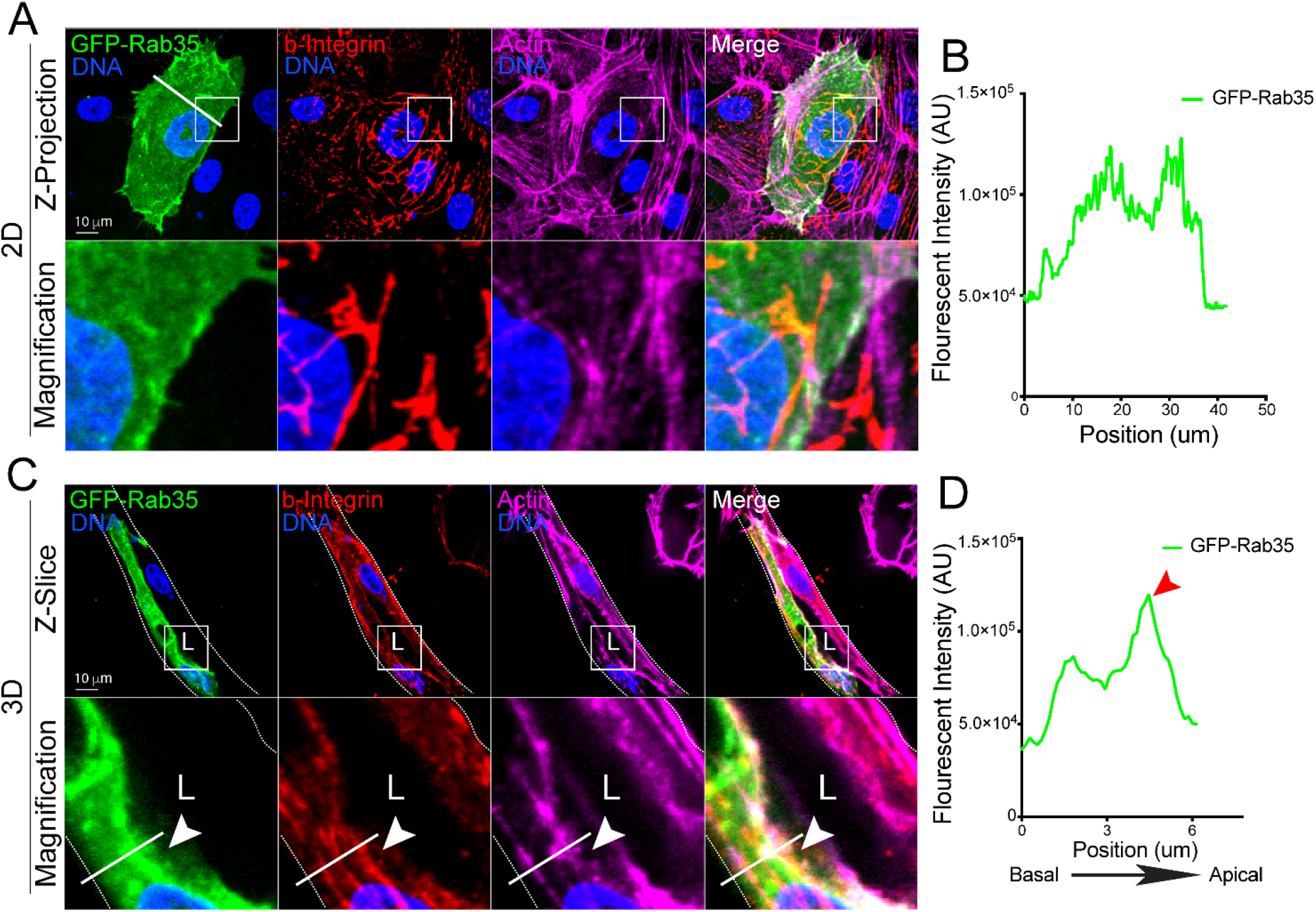
Visualizing Rab35 GTPase localization in 2D culture and 3D sprouts. (**A**) Representative image of endothelial cell expressing GFP-Rab35, stained for beta-1 integrin (b-integrin) and actin in 2D. Lower panels are magnification. (**B**) Line scan of Rab35 intensity in panel A. White line across cell in panel A represents line scan location. (**C**) Representative image of fibrin-bead sprout expressing GFP-Rab35, stained for b-integrin and actin. (**D**) Line scan of Rab35 intensity in panel C. White boxes are areas of magnification and while dotted lines indicated sprout boundaries. L denotes lumen.

Live-imaging in 2D culture showed GFP-Rab35 localized to dorsal membrane protrusions and lamellipodia; however, there was no membrane bias that was readily apparent (**Fig. 5A; Movie 3**). Live-imaging of Rab35 in the 3D sprouts revealed that, again, Rab35 is membranous with a preference to the apical membrane (**Fig. 5B**). Additionally, unique to the 3D sprouting environment, we could observe individual endosome movements adjacent to the apical membrane, which we could not detect in 2D culture (**Fig. 5B; Movie 4**). Interestingly, compared with 2D culture, the membrane dynamics were vastly dampened in 3D sprouting, suggesting culture in 2D may increase membrane dynamics consistent with other reports [40]. Overall, this data not only demonstrates a clear difference between Rab35 trafficking in 2D and 3D culture systems, but shows the importance of fully resolving apical and basal domains to interrogate function.

**Figure 5.**
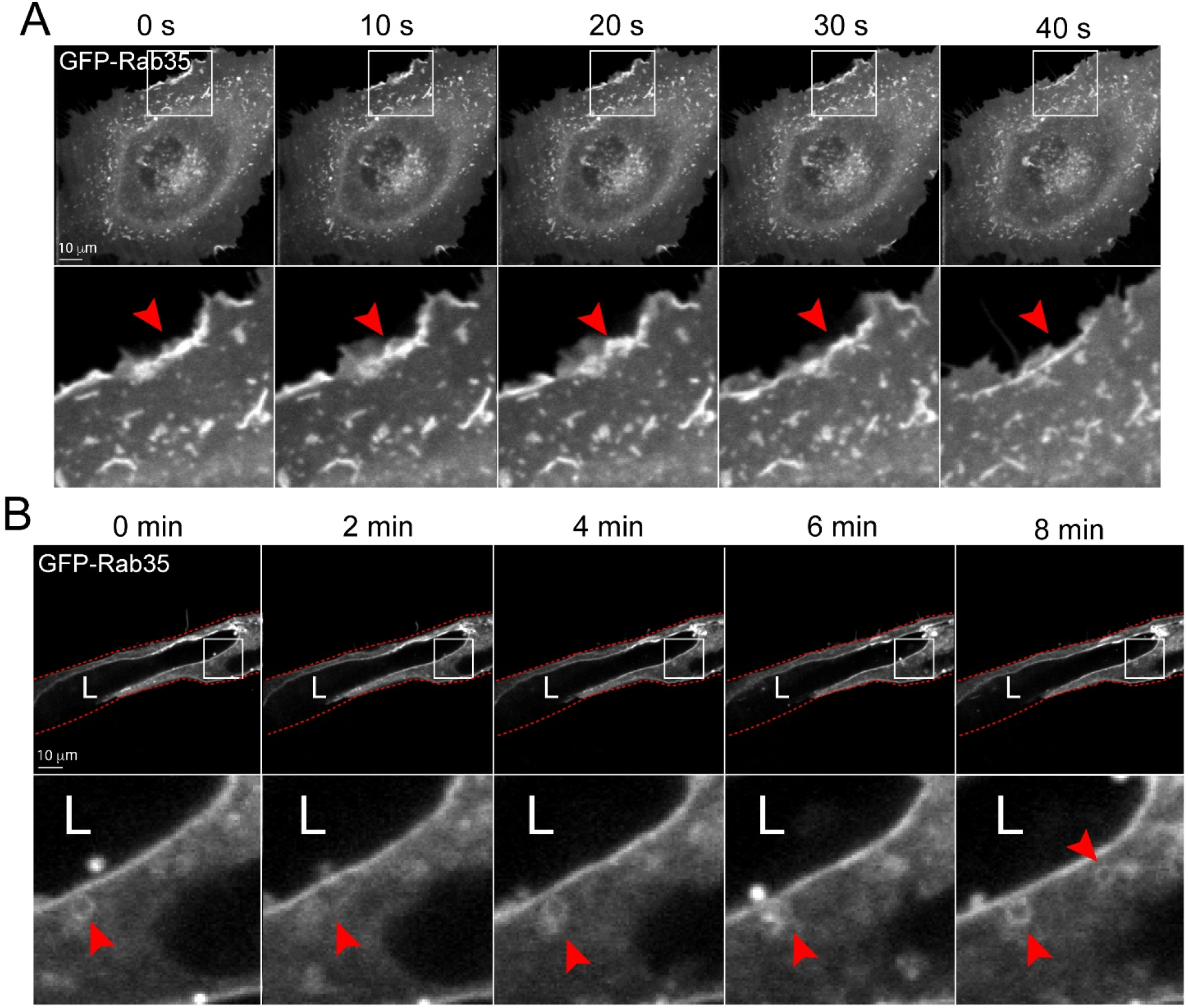
Live-imaging of Rab35 trafficking in 2D culture and 3D sprouts. (**A**) Live-imaging of cell expressing GFP-Rab35 over time. Red arrowheads denote puncta accumulated at the leading edge of the cell. (**B**) Live-imaging of fibrin-bead generated sprout expressing GFP-Rab35 over time. Red arrowheads indicate small endosome movements. White boxes are areas of magnification and red lines indicated sprout boundaries. L denotes lumen.

### Capturing exocytic events between 2D culture and 3D sprouting

An endothelial-specific trafficking function is the secretion of clotting proteins such as von Willebrand Factor into the luminal space upon injury [41-43]. WPBs are cigar shaped vesicles that are rapidly deployed to the apical membrane to exocytose many components including pro-thrombotic vWF [43]. We sought to determine if tracking WPB-related secretion events in 3D sprouts was feasible. We believe this is important as the majority of reports examining WPB biology use 2D culture opposed to tracking exocytic events in 3D sprouts with a more physiological defined apicobasal polarity. To do so, in 2D culture, we expressed GFP-Rab27a which has been previously shown to decorate WPBs [44-46]. Consistent with other reports Rab27a strongly co-localized with vWF on WPB puncta [44-46]. Co-labeled WPB puncta were randomly distributed throughout the cell with no distinguishable membrane preference (**Fig. 6A**). In 3D fibrin-bead sprouts Rab27a and vWF co-stained WPBs were easily visualized in proximity to apical or basal domains. In accordance with WPB function, we observed that WPBs accumulated at the apical membrane in many instances (**Fig. 6B**). Interestingly, we observed that WPB number is generally increased in ECs within sprout structures as compared with ECs in 2D culture (**Fig. 6C**). Of the WPBs in 2D culture there was almost equal split between the number of WPB accumulations that were in close proximity to the basal domain (≤ 1um) and generally contained within the body of cytoplasm, with 10% of WPBs at the apical domain (**Fig. 6D**). By contrast ECs in a 3D sprout showed that 36% of WPB accumulations were near the apical domain, 50% in the cytoplasm, 0% at the basal domain with the remaining 17% exocytosed into the lumen cavity (**Fig. 6D**). These results demonstrate that WPBs can be resolved 3D sprouts.

**Figure 6.**
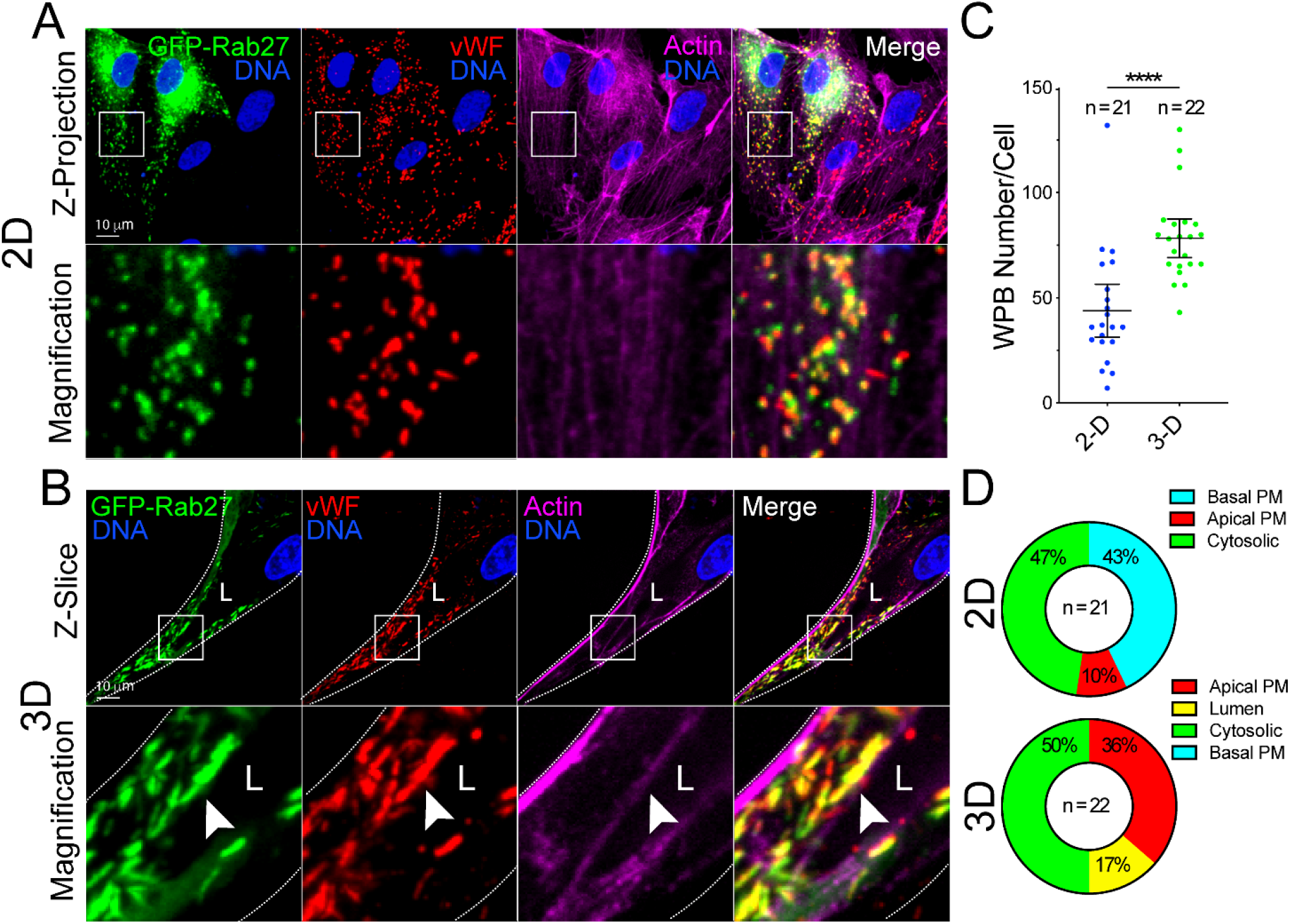
Imaging vWF exocytosis in 2D culture and 3D sprouts. (**A**) Representative images of endothelial cells in 2D culture expressing GFP-Rab27a and stained for von Willebrand Factor (vWF) and actin. (**B**) Representative images of a sprout expressing GFP-Rab27a and stained for vWF) and actin. White arrowhead denotes Weibel Palade Body accumulation. (**C**) Quantification of the number of vWF puncta between 2D culture and 3D sprouts. (**D**) Percentage of vWF accumulations (≥4 vWF puncta) that localize to a particular cellular location. PM= plasma membrane. N=number of cells. White boxes are areas of magnification and white dotted lines indicated sprout boundaries. L denotes lumen. Values are means +/- SEM; significance: ****P<0.0001. Statistical significance was assessed with an unpaired Students t-test.

Moving to live-imaging of WPBs labeled with Rab27a in 2D, we easily resolve discrete puncta and the disappearance of individual WPBs, presumably being exocytosed (**Fig. 7A; Movie 5**). In live-imaged 3D sprouts prior to lumen formation, we observed a robust accumulation of WPBs at the cell-cell interface as previously described by our group (**Fig. 7B; Movie 6**) [44]. Unlike 2D culture, we resolved WPBs aggregated at the apical membrane with less WPBs near the basal surface and were able to temporospatially track these structures relative to membrane domains (**Fig. 7B**). These results demonstrate that in the fibrin-bead sprouts sub-cellular WPB puncta can be tracked with enough resolution that monitoring movements between apical and basal surfaces can be achieved. This is an important aspect as exocytosis of WPB components will occur at the apical membrane.

**Figure 7.**
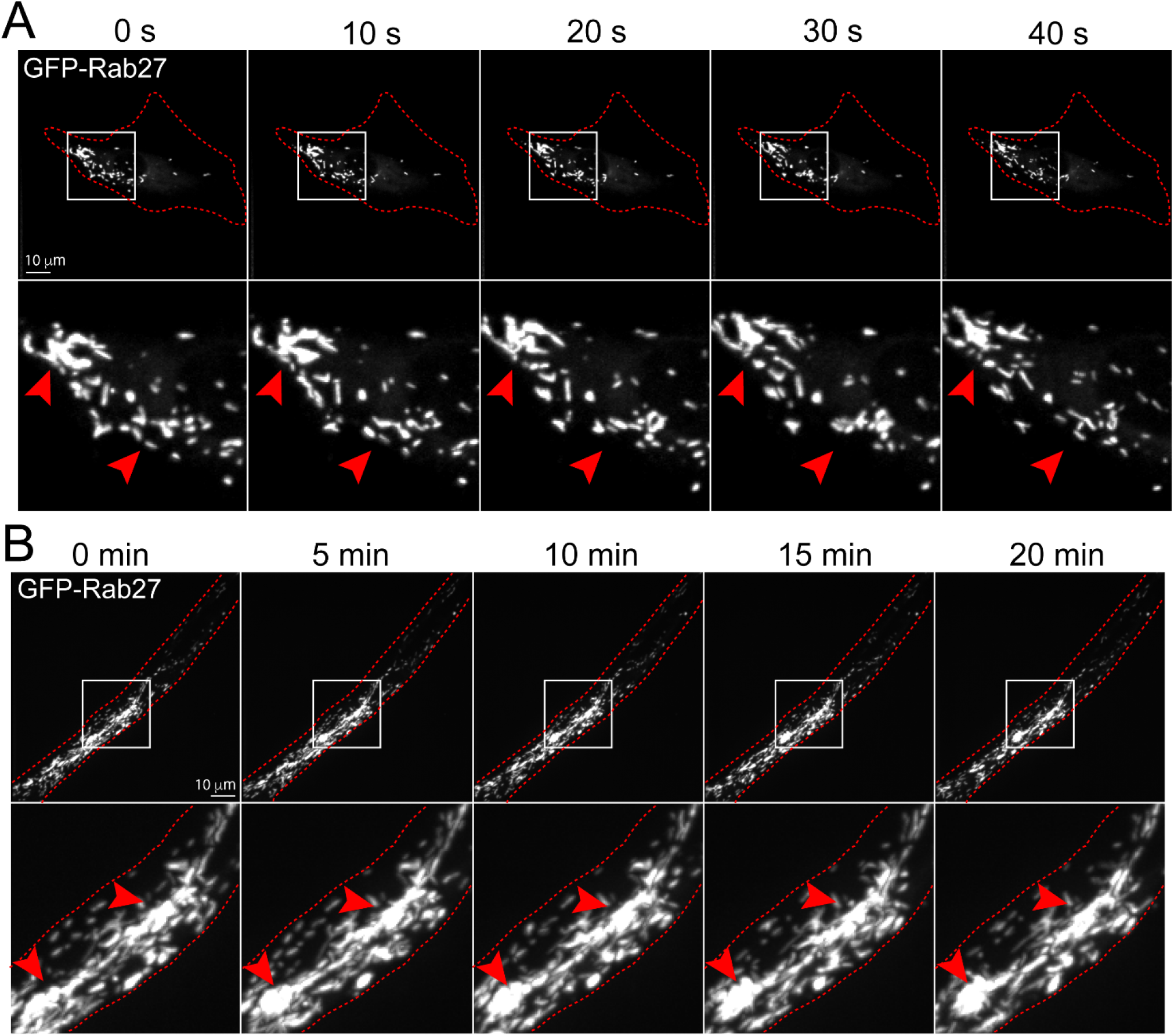
Live-imaging of Weibel Palade Body trafficking in 2D culture and 3D sprouting. (**A**) Live-imaging of endothelial cells expressing GFP-Rab27a (Weibel Palade Body marker) over time in 2D culture. Red arrowheads denote puncta accumulated at the leading edge of the cell. (**B**) Live-imaging of fibrin-bead generated sprout expressing GFP-Rab27a over time. Red arrowheads indicate accumulations of Weibel Palade Bodies at cell-cell interface. White boxes are areas of magnification and red botted lines indicated cell boundaries. L denotes lumen.

Next, we compared exocytosis of WPB-housed vWF between 2D culture and fibrin-bead sprouts. The rationale for this was: 1) the primary function of vWF is to be secreted, thus we wanted to capture this process in a 3D multicellular sprout structure; and 2) for a direct comparison to the standard 2D exocytosis assay of vWF many others have employed [10, 46, 47]. In 2D culture we knocked down Rab27a which has been shown to induce tonic vWF secretion [10]. In the Rab27a knockdown cells there was qualitatively less vWF puncta as compared with the scrambled controls; although, the amount of vWF exocytosis could not be assessed by imaging alone as it was secreted directly into the surrounding media (**Fig. 8A,C**). By contrast, Rab27a knockdown in fibrin-bead sprouts showed a dramatic accumulation of vWF effectively trapped within the sprout lumen (**Fig. 8B**). We could reproduce this result by administration of ionomycin (**Fig. 8D,E**), thus providing a conditional aspect to WPB evoked secretion. We believe this offers advantages over 2D culture in providing a better capacity to capture secretion events directed at the apical membrane in a growing sprout while simultaneously visualizing the relative amount of secreted protein(s) in the luminal space. These results show the utility of using a 3D sprouting culture system for visualizing exocytic events.

**Figure 8.**
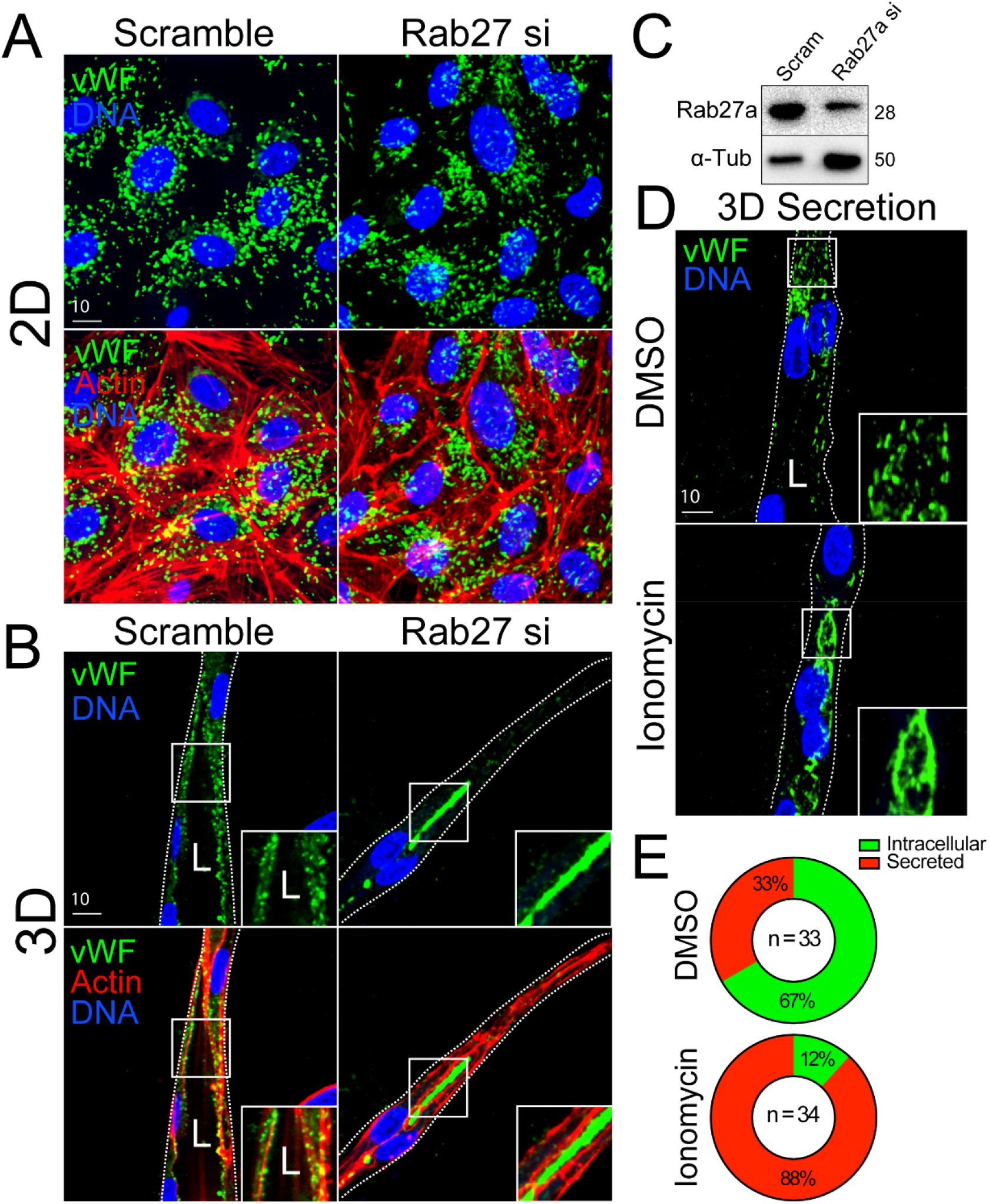
Monitoring exocytic events between 2D culture and 3D sprouting. (**A**) Representative 2D culture of endothelial cells stained for von Willebrand Factor (vWF) and actin treated with scramble or Rab27a-targeting siRNA (si). (**B**) Representative 3D fibrin-bead generated sprout stained for vWF and actin treated with scramble or Rab27a-targeting siRNA. (**C**) Western blot confirmation of Rab27a knockdown efficiency. (**D**) Representative 3D fibrin-bead generated sprout stained for vWF and treated with DMSO (vehicle) or ionomycin to induced Weibel Palade Body exocytosis. (**E**) Percentage of intracellular or lumen trapped vWF between indicated conditions. N=number of cells. White boxes are areas of magnification and white dotted lines indicated sprout boundaries. L denotes lumen.

## DISCUSSION

In the current investigation we show that the fibrin-bead sprouting assay has tremendous potential for imaging subcellular trafficking events. A primary disadvantage of *in vivo* imaging is the inability to capture trafficking processes due to the relative incompatibility of penetrating tissue with a long-working distance objective while maintaining high-enough resolution to resolve sub-cellular processes. Additionally, in many *in vivo* models, capturing dynamic events at the spatial scales required to distinguish trafficking processes, is not feasible. A common workaround to these issues is 2D culture of endothelial cells. However, our work establishes that this type of culture system strips away apicobasal signaling that is paramount to many trafficking processes. Our results using the fibrin-bead model highlights a middle ground where 3D sprouts that embody the most salient physiological characteristics of *in vivo* sprouting can be imaged at high-resolution on common confocal microscope platforms. Importantly, we demonstrate that imaging 3D sprouts not only presents advantages in capturing apical and basal domains due to sprout orientation, but is required to initiate proper apicobasal polarity. In the fibrin-bead we could clearly resolve trafficking mediators Rab35 and Rab27a both in live and fixed specimens and show their differential localization to the apical membrane. We also demonstrate the enhanced utility of the fibrin-bead assay for imaging exocytic proteins that are sequestered in the luminal cavity. Overall, our results support that the fibrin-bead sprouting model is suitable for visualizing endothelial trafficking events and could serve as an excellent companion assay to *in vivo* work.

Our data comparing fibrin-bead generated 3D sprouts with conventional 2D culture shows how trafficking, such as those in lumen biogenesis or secretion, are best imaged in 3D sprout structures. By virtue of being able to capture a cross-section of the sprout long-axis, imaging apical and basal domains is very accessible as this is the natural X-Y plane, unless acquired by lightsheet microscopy. Conversely, the apicobasal orthogonal view in cells imaged on a 2D surface needs to be digitally reconstructed limiting resolution. A problem that we originally envisioned was that the sprouts themselves would be too far from the coverslip limiting use of high numerical aperture, low working distance objectives. This was not the case, as most sprouts grew well within the working distance of a non-long working distance 60x objective. Static imaging of cell polarity markers provided a strong indication that virtually no apical signaling is present in 2D culture as compared with 3D sprouts as luminal proteins moesin and podocalyxin did not localize to the apical domain. Depending on the experimental question, this may not be a major issue. For instance, imaging of cytoskeletal proteins, this would likely not pose a problem. However, an issue arises when testing processes that require apicobasal polarity that is otherwise non-existent in 2D culture. This lack of polarity was also obvious when imaging Rab35 as it demonstrated apical localization in 3D sprouts and somewhat random localization on membrane protrusions in 2D cells.

A surprising finding was that exocytosis of vWF was perfectly contained within the luminal cavity when using the fibrin-bead assay. Imaging individual sprouts, we could very easily resolve discrete WPBs adjacent to the apical membrane, but could also potentially quantify the relative exocytosis of proteins on a per sprout basis given the secreted proteins are confined to the lumen cavity. In 2D culture, secreted proteins diffuse into the surrounding media blocking the ability to ascribe local secretion events to a particular group of cells. Given the preponderance of literature investigating WPB trafficking function using 2D culture, the fibrin-bead assay may be very beneficial in providing a more physiological component to these studies.

Culturing endothelial cells in 3D matrixes to induce sprouting behaviors is not new to the field of angiogenesis. Assays such as the Matrigel [21], vasculogenic assay [30, 48, 49], hanging-drop [50, 51], and fibrin-bead methods were initially used to investigate differences in endothelial sprouting and branching characteristics. For these types of gross morphometric analyses, these assays are still widely used; although, there are major advantages and disadvantages to each method. What has changed in more recent years is a focus on cell autonomous molecular mechanisms of sprouting angiogenesis that require an ever expanding need to visualize sub-cellular processes. In this regard, our group heavily employs the fibrin-bead assay, because as we focused on trafficking processes, we found a tremendous difference in 3D sprouting vs 2D culture systems. The differences in signaling between 2D plated cells and 3D sprouts was extreme enough to mask group changes in studies using loss and gain of function approaches. Not detecting differences makes sense if the 2D culture system does not provide apicobasal signaling cues, then perturbations in this polarity axis will not be readily apparent and lost to the investigator.

Our endorsement of the fibrin-bead sprouting assay does not exclude the possibility of other *in vitro* angiogenic or vasculogenic assays for imaging trafficking events as well as some *in vivo* models. For example, Davis et al. has demonstrated the excellent utility of the vasculogenic endothelial cell assay for not only tracking sprouting parameters, but visualizing sub-cellular processes like cytoskeletal proteins and caveolin localization [48, 52, 53]. Likewise, the original group who created the fibrin-bead assay has also engineered a microfluidic sprouting system that could be very interesting for investigating trafficking patterns with flow [27]. For *in vivo* models, unlike mammalians, zebrafish are optically transparent early in development allowing for live-imaging of blood vessel processes [54-57]. Many sub-cellular structures in endothelial cells can be distinguished using this model; however, in our hands, we cannot yield the resolution required to confidently quantify subtle trafficking localization events using needed 20x and 40x long working distance objectives. Therefore, we believe having the fibrin-bead assay to complement an *in vivo* model can help bridge deficits in either approach. There are many others who have developed comparable blood vessel sprouting-related assays that may be suited for imaging trafficking events that we have not mentioned. Our primary aim in the current investigation was to report our overwhelmingly positive experience with the fibrin-bead assay for imaging trafficking-related programs during angiogenic sprouting.

In conclusion, our data demonstrates that the fibrin-bead sprouting assay is an excellent platform for imaging endothelial trafficking events, particularly those related to apical and basal domains. We believe this method is a good substitute for 2D culture on many levels, most notably for imaging endothelial-related trafficking behaviors. We believe that endothelial-specific trafficking signatures represent a novel level of regulation that significantly contributes to vascular form and function. Thus, tools that aid in characterizing these processes will allow researchers to answer novel questions related to endothelial biology.

## CONTRIBUTIONS

C.R.F performed all experiments. C.R.F and E.J.K wrote the manuscript.

## Abbreviations

(2D): 2-dimensional
(3D): 3-dimensional
(WPB): Weibel Palade Body
(vWF): Von Willebrand Factor

## Supplemental Materials

### MAJOR RESOURCE TABLE

| Reagent | Vendor | Catalog # |
| --- | --- | --- |
| Poly-D-lysine | ThermoFisher | A3890401 |
| Polyethyleneamine Branched (PEI) | Sigma-Aldrich | 408727 |
| Chloroquine Diphosphate Crystalline (CQ) | Sigma-Aldrich | C6628-25G |
| Endothelial Cell Growth Medium 2 | PromoCell | C-22011 |
| DMEM, High Glucose, with L-Glutamine | Genesee Scientific | 25-500 |
| GenClone Fetal Bovine Serum (FBS) | Genesee Scientific | 25-514 |
| Penicillin-Streptomycin 100X Solution | Genesee Scientific | P4333-100ML |
| DPBS, no Calcium, no Magnesium | ThermoFisher | 14190250 |
| Trypsin-EDTA, 0.25% 1X, phenol red | Genesee Scientific | 25-510 |
| Paraformaldehyde 20% Aqueous Sol. EM Grade | Electron Microscopy Sciences | 15713 |
| Dimethyl Sulfoxide (DMSO) | Sigma-Aldrich | D2650-5X10ML |
| Silencer™ Negative Control No. 1 siRNA | ThermoFisher | AM4611 |
| Rab27 siRNA | ThermoFisher | siRNA ID: s11695 |
| Cytodex Microcarrier Beads | Sigma-Aldrich | C3275-10G |
| Fibrinogen Type 1-S from Bovine Plasma | Sigma-Aldrich | F8630-1G |
| Thrombin from Bovine Plasma | Sigma-Aldrich | T7513-500UN |
| Aprotinin Protease Inhibitor | ThermoFisher | 78432 |
| NHLF | Lonza | CC-2512 |
| HEK 293-A | ThermoFisher | R70507 |

### ANTIBODIES

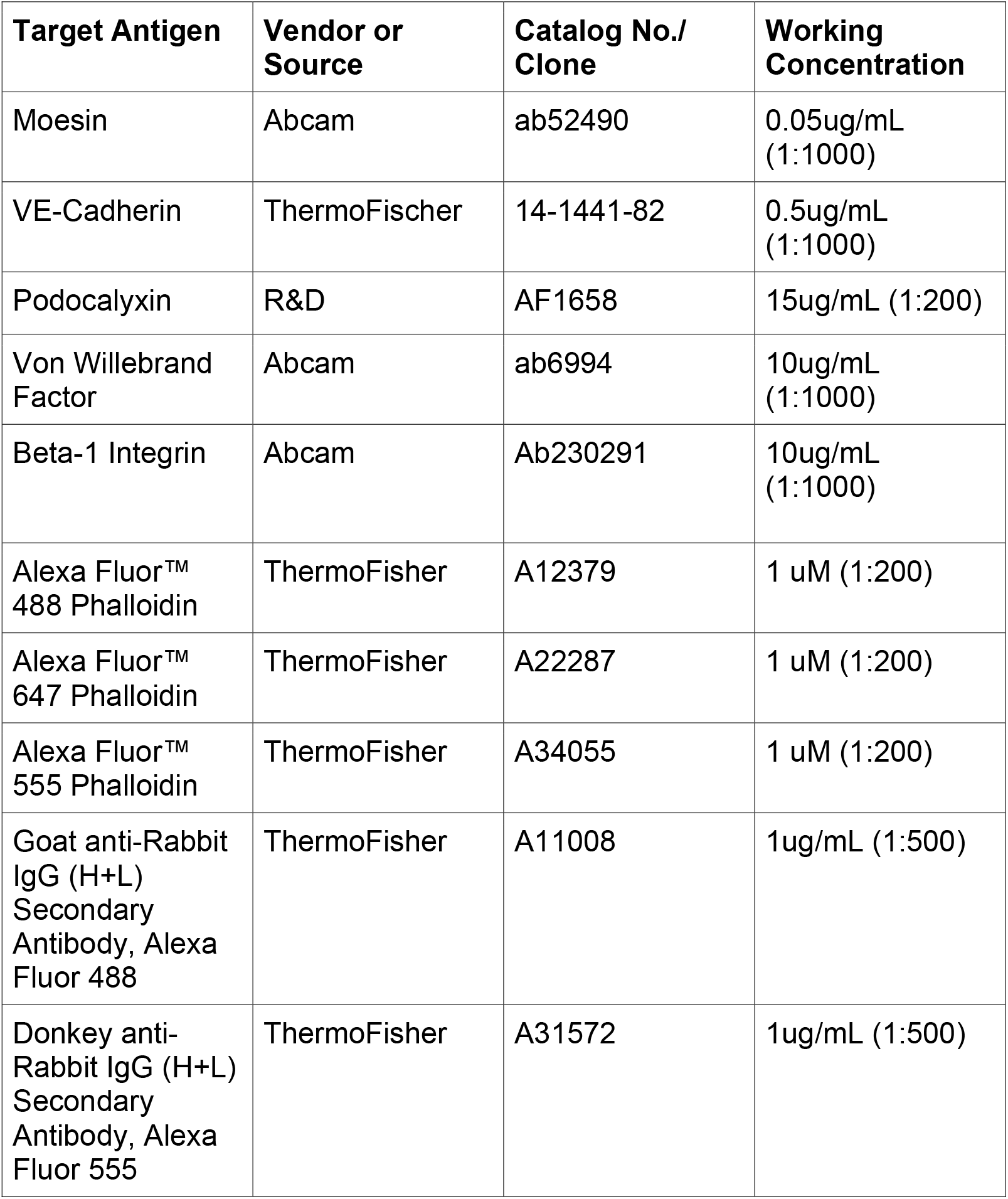
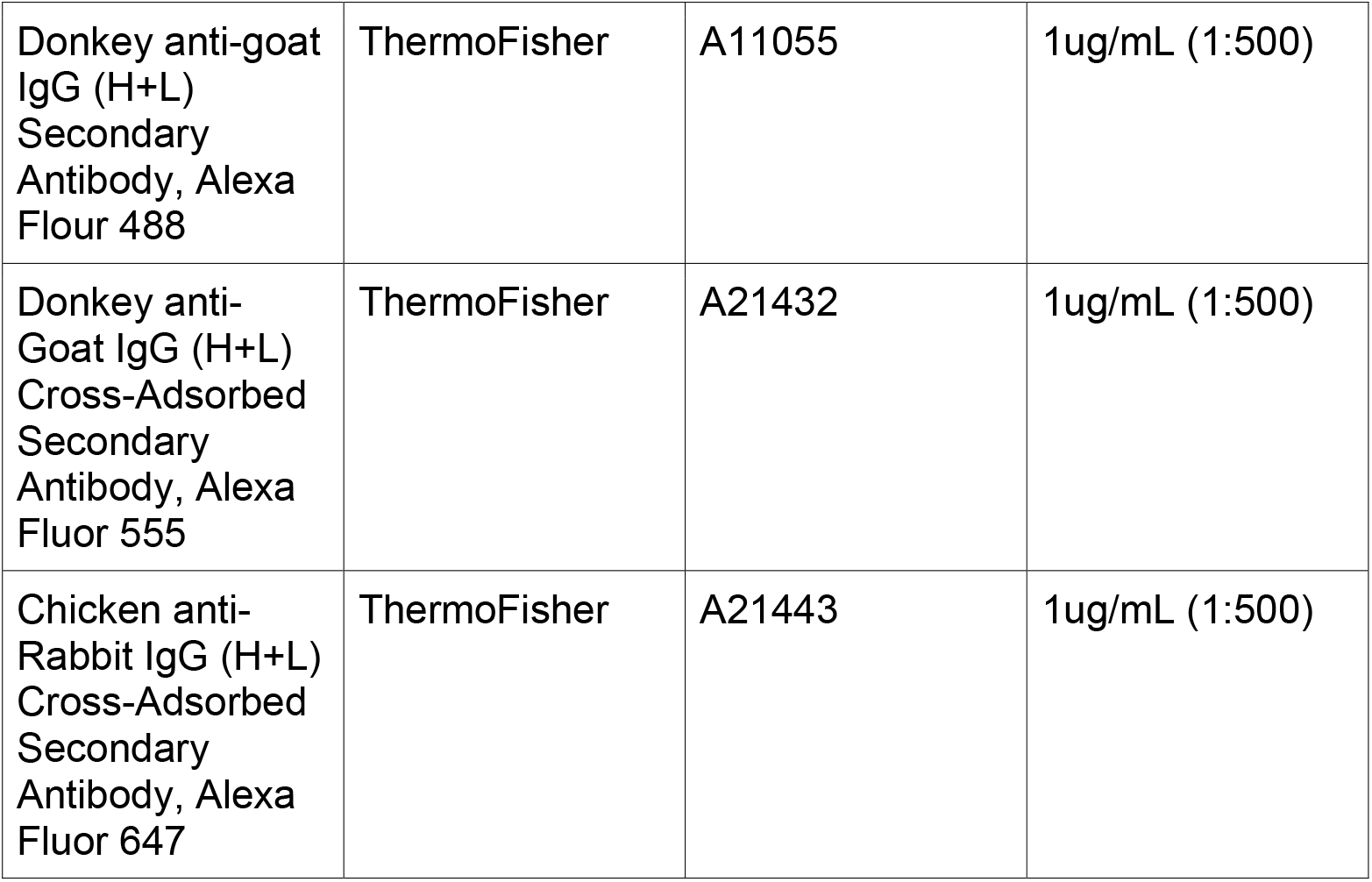

| Target Antigen | Vendor or Source | Catalog No./ Clone | Working Concentration |
| --- | --- | --- | --- |
| Moesin | Abcam | ab52490 | 0.05ug/mL (1:1000) |
| VE-Cadherin | ThermoFischer | 14-1441-82 | 0.5ug/mL (1:1000) |
| Podocalyxin | R&D | AF1658 | 15ug/mL (1:200) |
| Von Willebrand Factor | Abcam | ab6994 | 10ug/mL (1:1000) |
| Beta-1 Integrin | Abcam | Ab230291 | 10ug/mL (1:1000) |
| Alexa Fluor™ 488 Phalloidin | ThermoFisher | A12379 | 1 uM (1:200) |
| Alexa Fluor™ 647 Phalloidin | ThermoFisher | A22287 | 1 uM (1:200) |
| Alexa Fluor™ 555 Phalloidin | ThermoFisher | A34055 | 1 uM (1:200) |
| Goat anti-Rabbit IgG (H+L) Secondary Antibody, Alexa Fluor 488 | ThermoFisher | A11008 | 1ug/mL (1:500) |
| Donkey anti-Rabbit IgG (H+L) Secondary Antibody, Alexa Fluor 555 | ThermoFisher | A31572 | 1ug/mL (1:500) |
| Donkey anti-goat IgG (H+L) Secondary Antibody, Alexa Flour 488 | ThermoFisher | A11055 | 1ug/mL (1:500) |
| Donkey anti-Goat IgG (H+L) Cross-Adsorbed Secondary Antibody, Alexa Fluor 555 | ThermoFisher | A21432 | 1ug/mL (1:500) |
| Chicken anti-Rabbit IgG (H+L) Cross-Adsorbed Secondary Antibody, Alexa Fluor 647 | ThermoFisher | A21443 | 1ug/mL (1:500) |

## SUPPLEMENTAL MOVIES

**Movie 1. Live-imaging of pHluorin-podocalyxin in 2D culture**. Intervals are 1 frame/ second for 2 minutes.

**Movie 2. Live-imaging of pHluorin-podocalyxin in 3D sprouts**. Intervals are 1 frame/ 2 minutes for 20 minutes.

**Movie 3. Live-imaging of GFP-Rab35 in 2D culture**. Intervals are 1 frame/ second for 2 minutes.

**Movie 4. Live-imaging of GFP-Rab35 in 3D sprouts**. Intervals are 1 frame/ 2 minutes for 20 minutes.

**Movie 5. Live-imaging of GFP-Rab27a in 2D culture**. Intervals are 1 frame/ minute for 10 minutes.

**Movie 6. Live-imaging of GFP-Rab27a in 2D sprouts**. Intervals are 1 frame/2minutes for 20 minutes.

